# Age-related kinematic performance must be considered during fast head-neck rotation target task in individuals aged from 8 to 85 years old

**DOI:** 10.1101/519488

**Authors:** Renaud Hage, Frédéric Dierick, Nathalie Roussel, Laurent Pitance, Christine Detrembleur

**Author notes:** Corresponding author : Renaud Hage, Université Catholique de Louvain – Institut de Recherche Expérimentale et Clinique – Laboratoire NMSK. Avenue Mounier 53, B1.53.07 – 1200 Brussels, Belgium.

## Abstract

**Rationale:** Kinematic behavior during fast cervical rotations is a useful parameter for assessing sensorimotor control performances in neck-pain patients. However, in asymptomatic individuals from children to seniors, the influence of age still needs to be explored.

**Aim and method:** We assessed the impact of age on sensorimotor control performance of the head-neck with execution time and kinematic variables (time of task, mean speed/acceleration/deceleration, overshoots, minimum/maximum speed) during standardized fast rotation target task using the DidRen Laser test. Eighty volunteers were stratified in four different age-groups: Children [8-14y]: n=16; Young Adults [18-35y]: n=29; Old Adults [36-64y]: n=18; Seniors [65-85y]: n=17.

**Results:** To perform the test, Children were slower compared to Young Adults (p<0.001) and Old Adults (p<0.001). It was also slower in Seniors compared to Young Adults (p<0.013). Mean speed was slower in Children and Seniors compared to Young Adults (p<0.001) and Old Adults (p<0.001). Mean acceleration was slower for Children compared to Young Adults (p<0.016) and Old Adults (p<0.015). Mean deceleration was slower for Children compared to Young Adults (p<0.001) and Old Adults (p<0.003).

**Conclusion:** The DidRen Laser test allows us to discriminate age-specific performances for mean speed, acceleration and deceleration. Seniors and Children needed to be slower to become as precise as Young and Old people, no difference was observed for overshoots which assesses accuracy of movement. Age must therefore be considered as a key parameter when analyzing execution time and kinematic results during DidRen Laser test.

## 1 Introduction

The lifetime prevalence of neck pain is almost 70%, including pre- and adolescent patients, and this increases with age up to the age of 60 years ^1,2^. Nevertheless, underlying causes of neck pain are poorly understood. Because head rotation is a regular movement performed during everyday activities, this movement represents one of the issues that are of interest in patients with neck pain ^3^. Interestingly, previous studies supported that fast-rotational movements of the head-neck complex may be applied in clinical assessment and/or therapeutic management in patients with idiopathic neck pain ^4–7^. In view of these reports, fast axial rotation of the neck represents a critical motion to assess.

A target test using the DidRen Laser can be applied in response to real visual targets to standardize a head rotation motor task about 30° ^8,9^. Firstly, this test is believed to rely on the integrity of neuro-musculoskeletal structures and of the sensorimotor control system ^10^ i.e. input from the visual, vestibular and proprioceptive systems, particularly the richly innervated cervical spine, which controls head and eye movement and postural stability ^10^. Secondly, the axial head rotation about 30° will preserve the motion from the strain of the passive system (joint capsules, facets, intervertebral disks and ligament) ^11^.

Target-directed head-neck complex movements are fundamental components of individuals’ daily activities and usually require high levels of motion speed and accuracy. Speed-accuracy trade-off could be considered as a “signature” of the decision process ^12^. This varies according to which motion behavior is emphasized: accuracy or speed ^13^ during, for instance, an aiming task that encompasses amplitude of the movement, the size and position of the target ^5,12–14^.

Lack of sensorimotor control of head-neck complex can be explained ^15,16^ by degeneration of vestibular, visual and neuromuscular functions due to aging or even neck pain in adults and due to immaturity of the central and peripheral nervous and musculoskeletal system in healthy children^17,18^.

Sensorimotor performance of the head-neck complex can be assessed through various motor tasks requiring complex sensory input and motor output functions ^6^. These include the cephalic repositioning test, which estimates the proprioceptive and vestibular systems without interference of the visuomotor system. By using a laser beam placed on the head, the ability of blindfolded participants to actively relocate the head in a neutral position is evaluated after an active rotation movement of the head-neck complex ^19^. In other settings, the cervical proprioceptive system is assessed without focusing on the vestibular system by not including a cervical motor output component of the head-neck complex rotational kinematics. During this test, rotation of the trunk without movement of the head is measured with a 3D Fastrak electronic goniometer, by analyzing the effect of the modified cephalic repositioning test with a “neck torsion test” ^20^.

For including both sensory and motor component, other tests such as the “virtual reality test” ^9^ and “The Fly” ^21^ use a tracking system placed above the head to respectively measure the accuracy of the ability to follow a virtual target (the fly). Finally, the “DidRen Laser” is a test based on a system developed by our team in the late 2000s. It includes also both sensory and motor component by using a laser beam placed on the head of the participant with the aim to induce fast, low amplitude, and accurate rotational movements of the head-neck complex in response to real visual targets placed in front of the participant ^8^. Sensorimotor performance is assessed using the time difference between two successful hits of the targets. This means that, in view of the participant’s performance of speed-accuracy trade-off, the shorter the time, the faster and the more accurately the task is accomplished ^12,13,22^. In 2009, we showed that the DidRen Laser was a simple and reliable device with a good reproducibility in asymptomatic and symptomatic adult individuals ^8^. Compared to the other tests, the DidRen Laser is very easy to implement in a clinical setting, but it assesses only temporal variables. This does not allow us to gain insight into sensorimotor performance adopted by individuals.

In a recent study by Sarig Bahat et al. (2016), modified kinematics of the head-neck complex were reported in asymptomatic people over 60 compared to young and middle-aged adults ^23^. This study did not include measurements in children, which we consider essential to provide a better understanding of head-neck complex kinematics and normative data across all stages of the lifespan and to ensure that appropriate age-matched comparisons to participants with neck pain can be made ^24^.

To complete our previous results, it now appears essential to assess on asymptomatic individuals from children to seniors’, in addition of the temporal variables, further detailed kinematic and accuracy analysis carried out by a motion capture system, such as the maximum/minimum rotational speed, the average speed, the acceleration, deceleration and overshoot.

The aim of this study was then to analyze the effect of age from 8 to 85 years old on kinematic and accuracy variables derived from rotational motion of head-neck complex during the execution of a sensorimotor test called DidRen Laser. Our starting hypothesis was that sensorimotor control related to age would translate into slower kinematic and less accuracy performance when performing fast and precise neck rotations.

## 2 Material and methods

### 2.1 Participants

Eighty asymptomatic volunteers (43 females, 37 males) took part in this study and were stratified by age in four groups, children [8-14y]: n=16; young adults [18-35y]: n=29; old adults [36-64y]: n=18; seniors [65-85y]: n=17. They were recruited from colleagues in Saint-Luc University hospital (Brussels, Belgium) and among the researchers’ acquaintances. All participants volunteering for the study were informed about the nature of the study. Written consent was obtained from the participants, or from the legal representatives of participating children, before the start of the study. The study was approved by the local ethics committee and conducted in accordance with the declaration of Helsinki.

Participants suffering from any neuromusculoskeletal, neurologic disorder, impaired comprehension, non-correctable visual impairment, deafness, dizziness or any vestibular disorders diagnosed that could influence the performance of head-neck complex rotation, were excluded. Further inclusion criteria were absence of neck pain episodes in the 6 months prior to the study and to ensure that we analyzed asymptomatic participants, a Neck Disability Index (NDI) score of less than or equal to 4% ^25,26^. Similarly, all participants were asked to complete a Visual Analogue Scale (VAS) to confirm the absence of pain on the testing day ^27^. The main characteristics of the participants are listed in Table 1.

**TABLE 1.** Anthropometric characteristics of the participants

| Subjects | Global<br>(n=80) | Ch<br>(n=16) | YA<br>(n=29) | OA<br>(n=18) | S<br>(n=17) |
| --- | --- | --- | --- | --- | --- |
| Age (years), mean±SD | 40.25±28 | 11±2 | 24±3 | 53±7 | 73±5 |
| Sex (*men/women*), n | 38/42 | 5/11 | 13/16 | 11/7 | 9/8 |
| BMI ($kg\ m^{-2}$), mean±SD | 22.19±4.4 | 17.31±1.99 | 22.9±2.99 | 24.33±4.24 | 24.48±3.25 |
| NDI (100), median [Q1-Q3] | 2 [0-4] | 0 [0-2] | 2[1-2] | 2 [1-4] | 2 [0-4] |
| VAS (0-10), median [Q1-Q3] | 0 [0-0] | 0 [0-0] | 0 [0-0] | 0 [0-0] | 0 [0-0.5] |
| Global= all age-groups, Ch= Children, YA= Young Adults, OA= Old Adults, S= Old Adults, S= Seniors, SD= Standard Deviation, BMI = Body Mass Index, NDI=Neck Disability Index |  |  |  |  |  |

### 2.2 Instrumentation

The DidRen Laser was used (see ^8^ for a detailed description of the system and its components). In brief, it is made up of three photosensitive sensors with photocells, a fine laser beam, and a computer equipped with customized software that calculates the time between two consecutive “hits” of the sensors and displays the results. A chair without armrests is placed at 90 cm from a vertical panel equipped with the 3 sensors; each arranged horizontally and located 52 cm part from each other. The position of the central sensor is just in front of the head of the participant and 2 peripheral sensors induce the participant to perform right and left rotations of maximum 30°.

Three reflective markers were placed on the participant's head (Figure 1), two on the shoulders (acromion), and one on the cervical spine (C7). Three-dimensional recording of the markers positions were carried out at a sampling frequency of 200 Hz during the DidRen Laser test ^8^, using an optoelectronic system with 8 infra-red cameras (ELITE-BTS, Milan, Italy). A kinematic model composed of 6 anatomical landmarks (Fig. 1) was used. It was adapted from Bulgheroni et al. (1998) and representing the head and trunk segments. The head segment was modeled as a first triangle from the position of the head vertex (Top. H), and left/right side edges (R.H and L.H, respectively) landmarks (at 18 cm either side of the head vertex), and the trunk segment as a second triangle with left/right acromioclavicular joints (L.A and R.A, respectively) and the spinous process of C7 vertebra ^28^.

**FIGURE 1.**
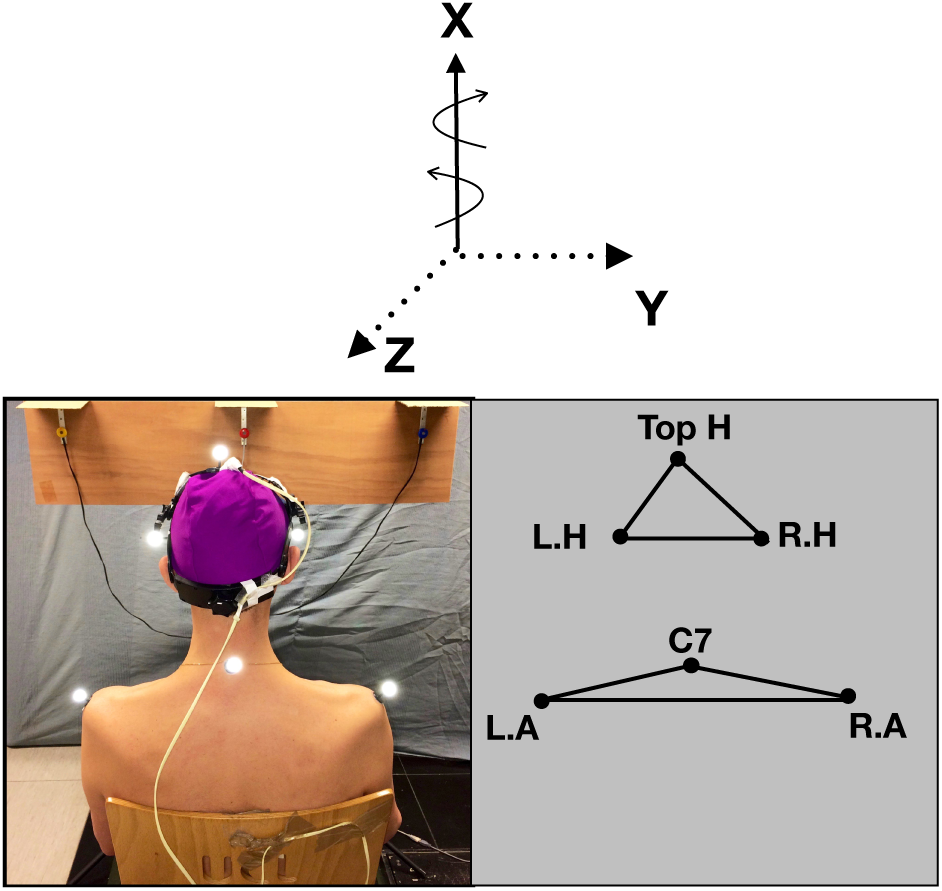
Head rotation is executed around a vertical axis (X) in a horizontal plane (Y-Z).

### 2.3 Experimental procedure

All participants received the same instructions about how the experimental procedure was to be conducted and more specifically that of the DidRen Laser test, by watching an explanatory video on a tablet computer. The participants wore a helmet to which a laser beam is mounted and sat on a chair. They were instructed to keep their back against the backrest, the palm of hands on the thighs, feet flat on the floor with heels against a stop block placed at the feet of the chair and to refrain from talking during the test.

The procedure of one generated cycle was explained as followed: turn your head faster as you can and point the laser correctly at the target. When the laser beam is pointed correctly at the target (during at least 0.5 s to lower chance), the LED’s sensor lights up and the system emits a sound signal. As soon as the central sensor has been ‘‘hit’’, the participant must, as rapidly as possible, turn his/her head to maximum 30° to the right to hit the right-hand sensor. He/she returns to the central sensor and then rotates the head to maximum 30° to the left to hit the left-hand sensor. The participant completes the cycle by hitting the central sensor for a third time. Then, one cycle is defined systematically with 4 head rotations facing sensors in the same order: 1) from center to the right, 2) from right back to the center, 3) from center to the left and 4) back to the center. A complete trial is composed of 5 cycles. Therefore, left and right rotation toward the target is composed by two phases. One fast rotation phase to turn the head followed by one stabilization phase to adjust the laser accurately during 0.5 second in the sensor/target (Fig.2). We used non-random generated cycles in order to facilitate intra and inter-subject comparisons.

**FIGURE 2.**
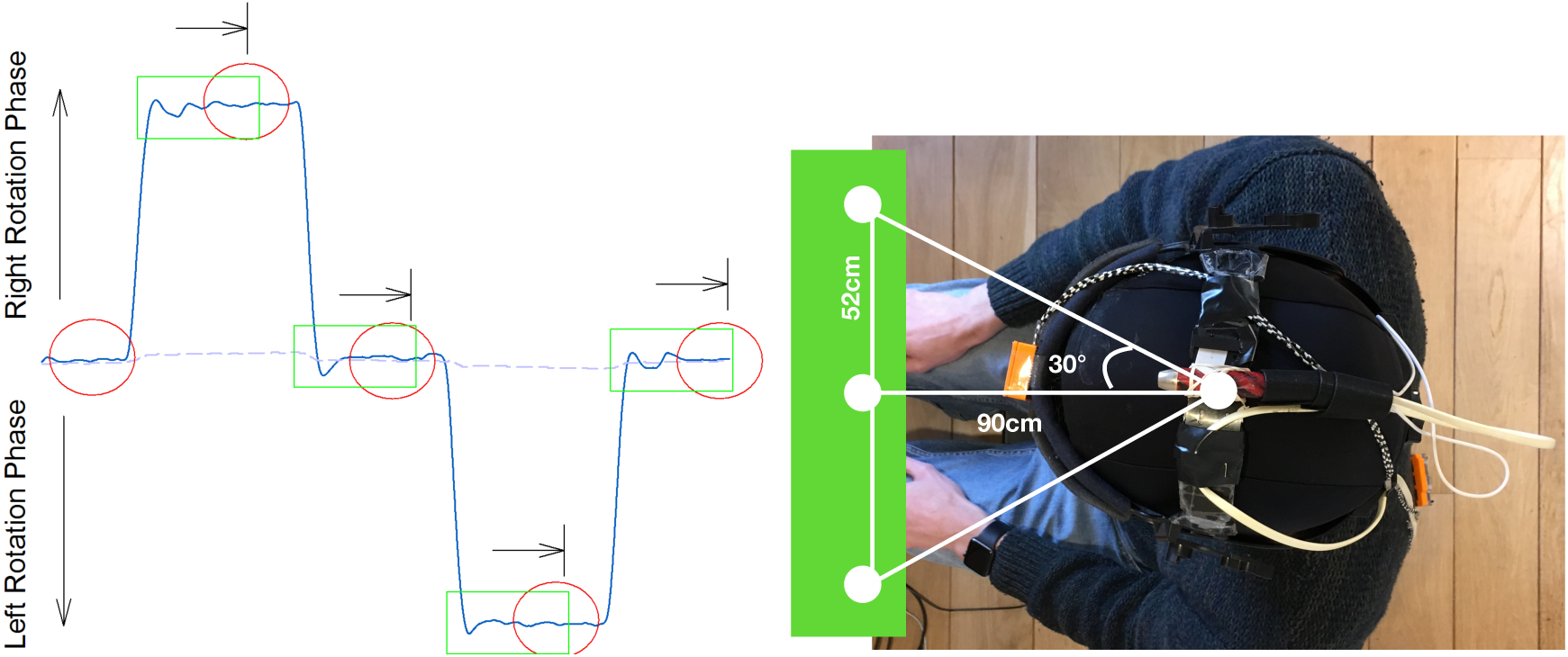
On the right: DidRen Laser installation device including the 3 photosensitive sensors. On the left: Typical trace of one rotational kinematic cycle showing Angular displacement versus time. Head rotations phases are visualized by the ascendant/descendant phase of the trace and stabilization phases are visualized by the horizontal phase of the trace. Circles depict targets.

As in 2009 ^8^, two DidRen Laser tests were conducted. The first trial was considered as a short warm-up to “familiarize” the participant with the experiment by emphasizing on the adequate sitting position and speed-accuracy execution. So only the second trial was used for data collection and analysis. We did not conduct more than two trials to avoid possible fatigue which could lead to a lost precision ^29^.

### 2.4 Outcomes measures

The DidRen Laser software calculates each time taken by the participant to go from one target’s ‘‘hit’’ (that is when participant stops during at least 0.5 s on the sensor) to another target. Only the total time (TT, in s) to complete the 5 cycles (from the first to the last target) of a trial was included in the data collection.

Head rotational and shoulders-displacement were computed using ELICLINIC software (BTS, Italy) from X, Y, and Z coordinates at each frame (Fig.1). To assess that the participants respected our theoretical calculation of 30° rotation without moving their shoulders, we calculated head-rotation range of motion (in °) and shoulders displacement (in °) (Fig.3). By successive numeric finite difference (n=±5 points), we calculated speed and acceleration of head. Because of their reliability, sensitivity and specificity ^4,23^, the following specific parameters were computed for each kinematics variable: maximum/minimum rotational speed (Max S, Min S, in °s^-1^); average rotational speed (Mean S, in °s^-1^) and average rotational acceleration/deceleration (Acc, Dec, in °s^-2^). To assess the accuracy, we calculated overshoot (OS, in °) (Fig.3) ^30^. It was computed as the difference between peak rotation amplitude and stabilized mean rotation amplitude. All variables were calculated during 5 consecutive cycles and averaged.

**FIGURE 3.**
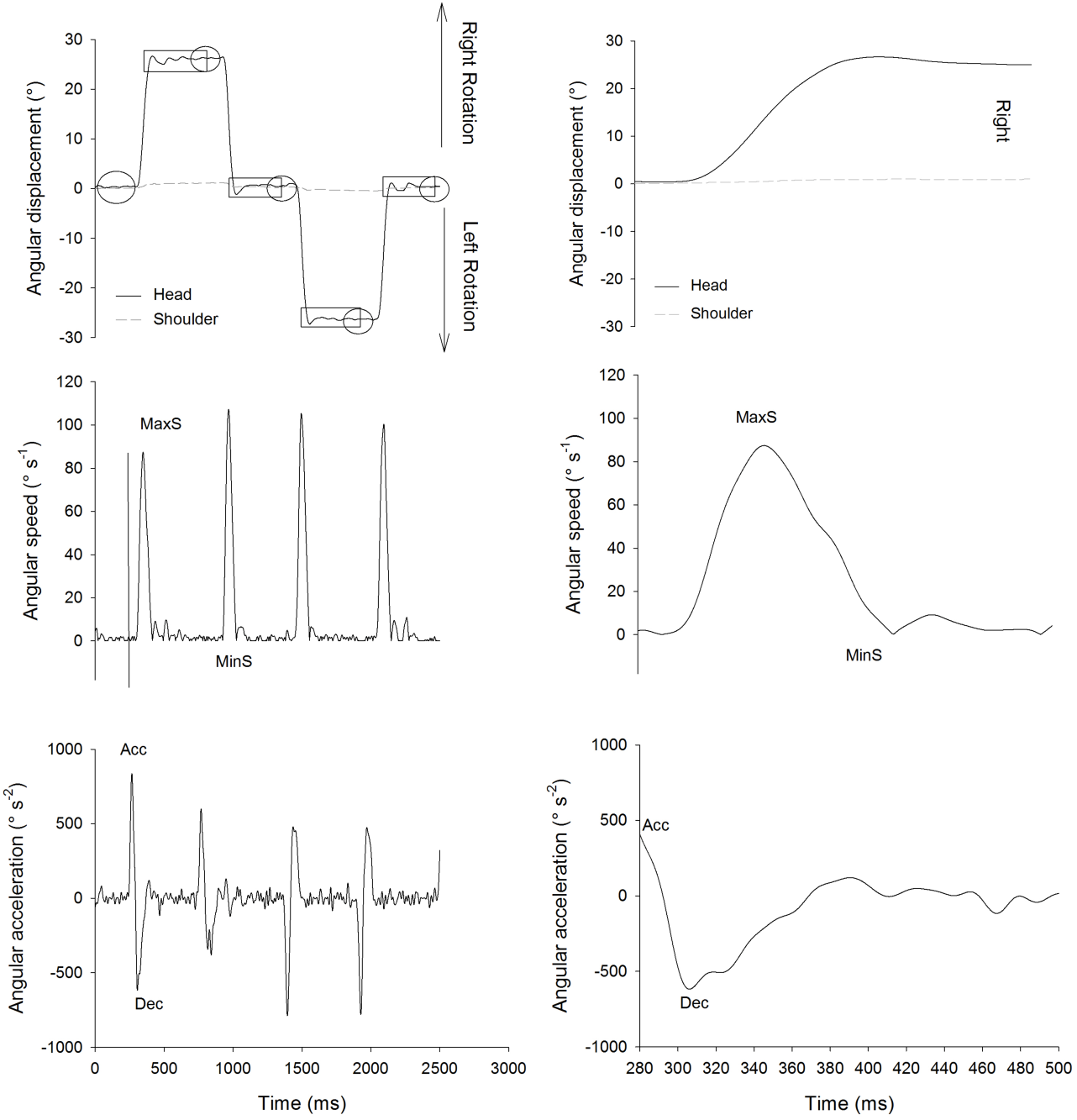
On the right: example of one rotational kinematic cycle with Angular displacement (showing Overshoot (OS) on rectangles and targets on circles), Angular speed (showing maximum speed (MaxS) and minimum speed (MinS) in absolute value) and Angular acceleration (showing acceleration (Acc) and deceleration (Dec)). On the left: example of one right rotation example during one cycle with Angular displacement, Angular speed (showing maximum speed (MaxS) and minimum speed (MinS) in absolute value) and Angular acceleration (showing acceleration (Acc) and deceleration (Dec)).

### 2.5 Statistical analysis

To assess the effect of age on the average variables, a one-way ANOVA with *post hoc* Holm-Sidak method for pairwise multiple comparisons was carried out when normally distributed and with *post hoc* Dunn’s method for pairwise multiple comparisons if normality test failed. All statistical procedures were performed with SigmaPlot 13 (Systat Software, Inc) with a significant determined at a of p<0.05.

## 3 Results

Total sample size consisted of 87 participants from which 7 participants were excluded due to an NDI score >4%. The anthropometric characteristics of the participants are listed in Table 1. Range of head-rotations did not exceed 30° (25.8±0.28°) and shoulders-displacements were negligible (0.7±1°).

All results are showed in Table 2 and Figure 4. A significant effect (p<0.05) of age was observed for 4 kinematic variables in Children and Seniors. The Total Time (TT (s)) was longer in Children compared to Young Adults (p<0.001) and Old Adults (p<0.001). It was also longer in Seniors compared to Young Adults (p<0.013). The average rotational speed (Mean S (° s^-1^)) was slower in Children and Seniors compared to Young Adults (p<0.001) and Old Adults (p<0.001). Acceleration (Acc (°s^-2^)) was slower for Children compared to Young Adults (p<0.016) and Old Adults (p<0.015). Deceleration (Dec (°s^-2^)) was lower for Children compared to Young Adults (p<0.001) and Old Adults (p<0.003).

**Fig. 4.**
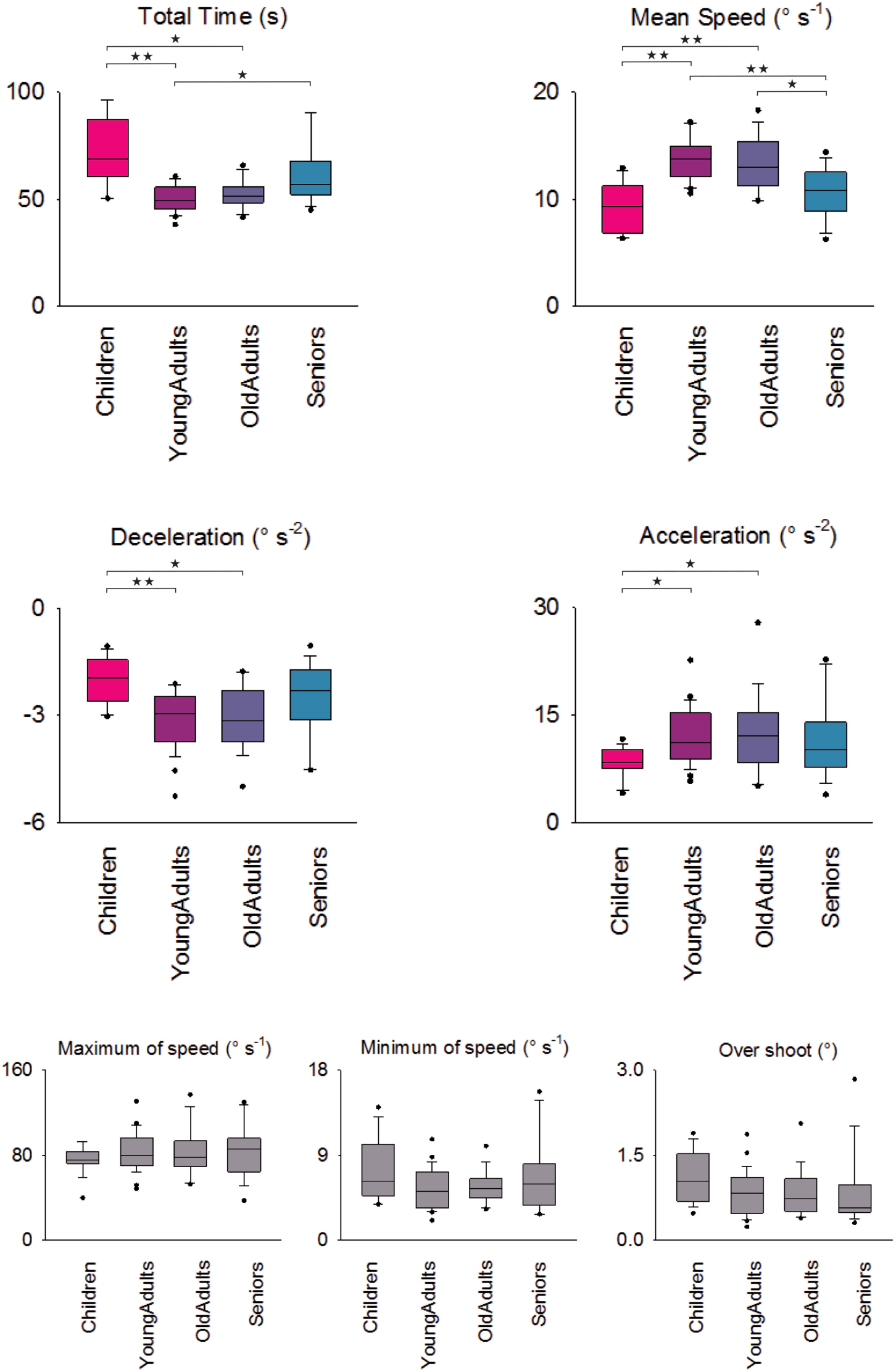
Box-plots (interquartile range) for all age effect measurements. First four box-plots are for age-groups significant kinematic variables with *post hoc* comparisons: *★ P<0.05, ★★ P<0.001*. The three horizontal box-plots are for non-significant kinematic variables.

**TABLE 2.** For all variables, if normality test passed results were in mean with standard deviation (SD) and if normality test failed results were in median with interquartile range [Q1-Q3]

| Variable | Tests | Ch | YA | OA | S | Comparisons between groups |
| --- | --- | --- | --- | --- | --- | --- |
| **<u>TT(s)</u>** | Mean [Q1-Q3] | 69.07 [60.657.4-879.34] | 49.6 [45.6-55.6] | 51.7 [48.4-55.8] | 57 [52.3-67.6] | Ch > YA**, OA*<br>S > YA* |
| **<u>Mean S (°s<sup>-1</sup>)</u>** | Mean±SD | 9.4 ± 2.3 | 13.7 ± 1.9 | 13.3 ± 2.4 | 10.6 ± 2.4 | Ch < YA**, OA**<br>S < YA**, OA* |
| **<u>Dec (°s<sup>-2</sup>)</u>** | Mean [Q1-Q3] | -1.9 [-2.6--1.4] | -2.9 [-3.7—2.5] | -3.2 [-3.7—2.3] | -2.3 [-3.1--1.7] | Ch > YA**, OA* |
| **<u>Acc (°s<sup>-2</sup>)</u>** | Mean [Q1-Q3] | 8.4 [7.6-10.2] | 11.1 [8.8-15.3] | 12.0 [8.4 -15.3] | 10.2 [7.7-14.0] | Ch < YA*, OA* |
| **<u>Max S</u>** | Mean [Q1-Q3] | 75.1 [72.2-83.2] | 79.9 [70.1-96.3] | 77.7 [69.1-93.0] | 85.8 [64.2-95.6] | # |
| **<u>Min S (s)</u>** | Mean [Q1-Q3] | 6.2 [4.7-10.1] | 5.2 [3.4-7.2] | 5.4 [4.5-6.5] | 5.9 [3.6-8.0] | # |
| **<u>OS (°)</u>** | Mean [Q1-Q3] | 1.0 [0.7-1.5] | 0.8 [0.5-1.1] | 0.7 [0.5-1.1] | 0.5 [0.5-0.9] | # |

Ch=Children, YA= Younger adults, OA= Older adults, S= Seniors. Age-related significant difference is observed for time duration (DidRen total time), for Mean Speed (MeanS), Deceleration (Dec) and Acceleration (Acc). No age-group significant difference = (#) for Maximum speed (Max S), Minimum speed (Min S) and Overshoot (OS). Indications for which group differed from another: Longer or Slower = (>), Less = (<). \* P<0.05 \*\* P<0.001

## 4 Discussion

The aim of this study was to analyze the effect of age from 8 to 85 years old on sensorimotor control performance adopted by asymptomatic individuals using kinematic and accuracy variables derived from rotational motion of head-neck complex during the execution of the DidRen Laser test.

The results of the study reveal significant effects in asymptomatic Children and Seniors groups for four rotational kinematic variables: TT, Mean S, Acc and Dec. These kinematic differences were observed during the DidRen Laser test, showing its capacity to discriminate age-related differences. Children-age limit of 14 years was based on need to ensure of an incomplete maturation of the sensorimotor representation ^17^. Moreover, we decided to split adults participants into two adults groups because neck pain increase with age and is most common around the fifth decade of life ^1^.

Velocity and its time derivate forms were chosen to analyze the performance of age-group participants in accordance with previous studies ^4,9^. Sarig Bahat et al. (2016) showed that the most powerful age-groups differences were Velocity Peaks and Number of Velocity Peaks ^23^. With a different task and method of calculation, our kinematic variables such as TT, Mean S, Acc and Dec appeared to be very significant especially with the groups of Children and Seniors.

To reach the speed-accuracy required by our protocol: “turn your head faster as you can and point the laser beam correctly at the target”, participants needed to trade-off with their sensorimotor ability to perform first the fast-rotational movement and then to enable the participant to stabilize the motion and to adjust the laser into the sensor of the target ^12^. The task reflects an orchestrated pattern of neural activation to select a minimize time to reach target ^31^, but participants needed to trade accuracy for speed because they were instructed to respond as fast as possible to a constrained target-aiming task ^13,16,32^. In light of the foregoing, we showed that Seniors and Children needed to be slower (TT, Mean S, Acc, Dec) ^5,14,32,33^ to become as precise as Young and Old people (no significant differences for OS in all age-groups). The validity and reliability of the OS seems to be good when comparing patients with WAD with controls ^21^ and with patients with non-traumatic neck pain ^30^. Even if we have calculated OS differently than Kristjansson et al. (2004), this concept of analyzed is representative of the quality of the cervical sensorimotor status. Our results have then demonstrated the ability for Children and Seniors to adapt their speed to be at least as accurate as Young and Old Adults. Similarly, we have also demonstrated that both Adults groups have adapted their speed. Indeed, both Adults groups showed an average of speed 10 times less than the average of speed (Vmean) obtained by Sarig Bahat’s participants of the same ages ^23^. Actualy, even if Sarig Bahat’s participants were asked to move their head fast, they could take longer time (seven seconds) to point the target after having produced the motion. So, they did not need to trade-off with their speed and accuracy as much as needed to accomplish the DidRen Laser test. However, notwithstanding with our empirical speed-accuracy trade-off observation, we can’t be sure that speed adaptation is not consequence of the mechanical properties of muscles which we know can influence coordination during functional activities like cervical rotation ^34^. Because older age does influence performance in muscles activity during a cranio-cervical flexion coordination test ^34–36^, this could lead to a reduction in the cortical representation of the stabilizing muscles and thus, hinder automatic contraction during head movement as rotation ^37^. Unfortunately, to our knowledge the influence of neck muscle activities during cranio-cervical flexion coordination test in children is not known.

The DidRen Laser test ^8^ offers the advantage to focus on the sensorimotor control system of the head-neck complex with many direct neurophysiological connections between the proprioceptive, the visual and the vestibular systems ^38^. It involves real visual targets completed by an auditory feedback system: when the laser beam is pointed correctly at the target, the sensor lights up and the system emits an audible sound signal. Even if the kinematic variables assessed in this study do not allow us to differentiate the subsystem(s) of the sensorimotor control, it is plausible that senior-related modifications can be due to dysfunction in neck proprioception ^15,39^, vestibular ^40^ and visual systems ^15^. Slower Children-related performances could be due to slow maturation of the sensorimotor system ^18,41^ and immaturity in the development of the central nervous system ^18,41–43^.

The results of the present study should be seen in the light of several methodological clarifications. Firstly, standardization of instructions and better understanding of participants, especially for children and some older participants, were increased by using an explanatory video of the DidRen Laser test. Secondly, to avoid measurement errors that could increase variability of our results, we standardized the posture of the head, trunk, limbs and the height of visual reference and the influence of the scapular girdle displacement while performing head-neck complex rotations ^44–46^. Thirdly, to avoid influencing stability of the neck and for being more functional, we did not secure the shoulder with a seat-belt ^47^. Fortunately, our results concerning the scapular girdle rotation displacement showed that it was marginal for all participants and confirmed that they respected the instruction not to move their shoulders during the test. Thanks to this option, the DidRen Laser test could be used more easily for future clinical studies.

This study presents some limitations. First, neck ROM was limited to 30° that allowed us to avoid the strain of the neck passive system (joint capsules, facet joints, intervertebral disks and ligaments) and to improve input from the upper cervical proprioceptive system which is highly developed in the sub-occipital region upper neck ^48^ and which corresponds to the spinal muscles that provides dynamic stability during the first degrees of rotation ^11^. Second, only head-neck rotational movement was assessed but rotation seems to be a regular movement during daily activities, the assessment of other motion directions (e.g. flexion/extension) appeared to have limited interest. Third, we used for the test regular generating head rotation cycles. We acknowledge that the use of randomly generated cycles might reduce an induced anticipatory motions and predictions of participants but that has not yet been investigated for the DidRen test.

In conclusion, DidRen Laser test allows us to discriminate age-specific performances for mean speed, acceleration and deceleration. Empirically we showed that Seniors and Children needed to be slower to become as precise as Young and Old people by showing no difference for overshoots which assesses accuracy of movement. Age must therefore be considered as a key parameter when analyzing execution time and kinematic results during DidRen Laser test but not for accuracy.

